# A simple method of mechanical scarification to break seed dormancy in *Viola odorata*

**DOI:** 10.1101/2025.11.05.686783

**Authors:** Joo Young Kim, Karina Idiyatullina, Thomas A. Colquhoun

**Affiliations:** Environmental Horticulture Department, Plant Innovation Center, Institute of Food and Agricultural Sciences, University of Florida, Gainesville, FL 32611, USA

**Author notes:** Corresponding author: Thomas A. Colquhoun, 1529 Fifield Hall, University of Florida, Gainesville, FL 32611, USA.

**Keywords:** cold stratification, gibberellic acid, water uptake, germination, sweet violet, *V. odorata* ‘Express Augusta’

## Abstract

*Viola odorata*, also known as sweet violet, is a valuable plant for its fragrance and medicinal properties, but the cultivation is hindered by poor seed germination due to dormancy. The dormancy of *V. odorata* mainly resulted from a hard seed coat, and various methods has been reported in breaking the seed dormancy. However, most engaged in corrosive chemicals and highly dependent on the variety, or a lack of uniformity when using sandpapers. This study presents a simple, effective mechanical scarification method using rat-tooth tweezers to gently nick the seed coat tip. This mechanical scarification significantly increased germination, reaching a 70% rate after eight weeks, while non-scarified seeds failed to germinate. Combining mechanical scarification with cold stratification further enhanced germination compared to room-temperature (RT) storage. Additionally, growth at 18 °C produced higher rates than at RT and the addition of gibberellic acid (GA_3_) in the media promoted both germination and seedling growth, accompanied by increased water uptake. These findings propose a combination of mechanical scarification, cold stratification, and optimal GA_3_ treatment as a reliable, broadly applicable strategy for breaking dormancy and facilitating large-scale propagation and breeding of *V. odorata*.

## Introduction

*Viola odorata*, commonly known as sweet violet, is a perennial species that belongs to the Violaceae family. Native to Europe and Asia, the plant has been naturalized in North America and Australasia (Marcussen, 2006). It is recognized for its strong fragrance and the presence of various bioactive compounds, including alkaloids, saponins, cyclotides, and flavonoids (Mittal et al., 2015). Research has highlighted the potential medicinal effects of *V. odorata*, such as anti-inflammatory, antipyretic, and antibacterial properties, as well as its ability to enhance liver function (Mittal et al., 2015). The viola’s therapeutic and ornamental value has also led to growing interest in research as a genetic resource for fragrance in plant breeding (Shea and Colquhoun, 2022).

Despite its significance, the propagation rate from *V. odorata* seeds is relatively low due to their dormancy. Dormancy is a critical adaptive mechanism that prevents seeds from germinating under unfavorable environmental conditions, thereby increasing the probability of successful regeneration and enduring the long-term persistence of the species (Nawrot-Chorabik et al., 2021). Baskin and Baskin (2004) categorized dormancy into several types: physiological, morphological, morphophysiological, physical, and combinational dormancy. *V. odorata* seeds exhibit non-deep physiological dormancy, which is attributed to their relatively hard and water-impermeable seed coats, particularly water-impermeable layers in their seed coats (Baskin and Baskin, 2004). The seed coats also act as physical barriers that inhibit the growth of the embryo. For germination to occur, the seed coat must become permeable to allow water uptake, and the embryo’s growth potential must be sufficient to overcome the mechanical resistance of the seed coat (Finch-Savage and Leubner-Metzger, 2006). A variety of methods are known to effectively break physiological dormancy, including mechanical scarification, cold and chemical stratification, controlled lighting conditions, and the application of plant hormones such as gibberellic acid (GA_3_) (Baskin and Baskin, 2004; Finch-Savage and Leubner-Metzger, 2006; Mousavi et al., 2011). GA_3_ is known for stimulating embryo growth and helping break seed dormancy by promoting the production of enzymes that mobilize stored nutrients and soften the seed coat (Gupta and Chakbarty, 2013).

Although significant research has been conducted on seed dormancy, there are few studies have focused on improving the germination of *V. odorata*. Keene and Colquhoun (2022) tested different concentrations of GA_3_ and the duration of cold stratification. The treatments significantly improved germination in *V. odorata* ‘Rubra’, particularly when seeds underwent stratification for 8 and 12 weeks at 4°C. Barekat et al. (2013) reported that embryo culture and treatment with concentrated sulfuric acid resulted in higher germination rates, while unstratified *V. odorata* seeds failed to germinate. This suggests that mechanical scarification is a highly efficient method for breaking the seed dormancy of *V. odorata*. However, these methods have some limitations. The use of concentrated sulfuric acid is highly corrosive and poses risks upon contact with skin, and the efficacy of GA and cold treatments can vary depending on the variety. While embryo culture is an effective approach for germinating *V. odorata* seeds, the complete removal of seed coats without causing damage is not easy. Here, we report a precise and non-corrosive mechanical scarification technique that improves germination and investigate additional conditions to increase the germination rate of viola seeds. Low germination rates in viola seeds can hinder large-scale propagation and breeding programs, thereby limiting the effectiveness of efforts to enhance genetic diversity (Keene and Colquhoun, 2022). This research benefits in improving seed germination, leading to genetic diversity in *V. odorata* propagation and breeding.

## Materials and Methods

### Plant materials and culture conditions

Seeds of various *Viola odorata* cultivars, including ‘Empress Augusta (EA)’ were purchased from Gardens in the Wood of Grassy Creek (Crumpler, NC, USA). Seeds were stored at either room temperature (RT) or 4 °C for a minimum of 3 months. For sterilization, seeds were washed with deionized (DI) water and soap, then soaked in DI water for one hour to soften the seed coats. After soaking, the tips of the seed coats were cut for mechanical scarification. The cut seeds were sterilized using the method described by Barekat et al. (2013). This involved soaking the seed in 70% ethanol for 1 min, followed by three rinses with sterile DI water. The seeds were then soaked in 1% sodium hypochlorite with Tween-20 (drop/mL) for 15 min, followed by four rinses with sterile DI water. Finally, the seeds were dried overnight in a clean flow hood.

The basal media for all experiments consisted of half-strength Murashige and Skoog (MS) medium with vitamins, supplemented with 15 g/L sucrose, 0.5 g/L of 4-morpholineethanesulfonic acid (MES), and 7% agar. The pH of the medium was adjusted to 5.7. Thirty sterilized seeds were placed onto each petri-dish containing the medium, with three replications per treatment. The petri-dishes were maintained in darkness at 18 °C in a plant growth chamber (Percival Scientific, Perry, IA, USA).

### Mechanical scarification

For mechanical scarification, the outer seed coat of EA was carefully cracked at the tip using rat-tooth tweezers (Figure 1A and 1B) to expose the internal structures, including the embryo tip, without causing damage (Figure 1C). Each seed was mechanically scarified by gently nicking the tips of the seed coat with rat-tooth tweezers to make a small incision. The cut was made just deep enough to slightly open the hard outer layer near the embryo without damaging it. The seeds were sure not crushed, and the embryo remained intact. This method was developed as a safer and simpler technique for seeds that require stratification to germinate.

**Figure 1.**
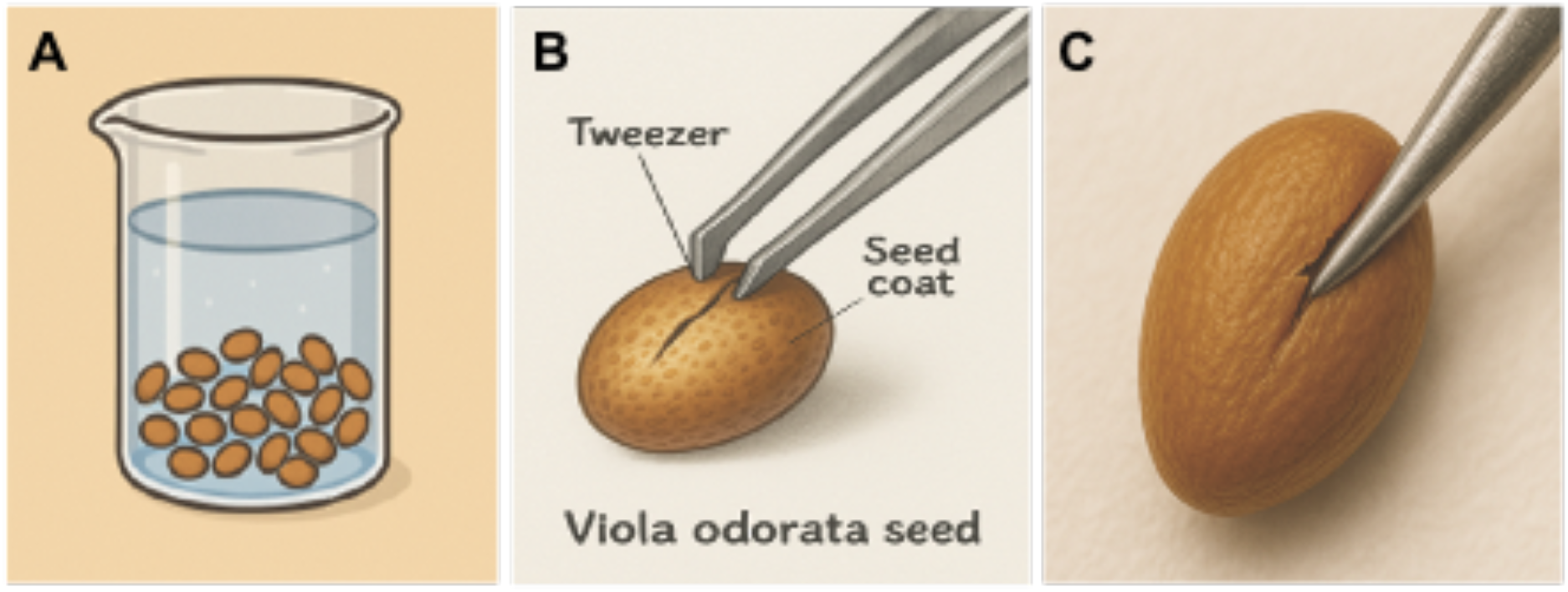
Easy method of mechanical scarification. Soak viola seeds in water for one hour (A) and cut the seed coat with a rat-tooth tweezer (B), resulting in opening for seeds without damaging the embryos (C).

### Experiment 1: Cold stratification and growth temperature

The effect of cold stratification on the germination of EA seeds was investigated. Seeds were stored at either RT or 4 °C for a minimum of three months. Mechanically scarified EA seeds were sterilized and placed on MS media and grown in a growth chamber maintained at 18 °C for 8 weeks. Germination rates of cold-stratified seeds were compared with those of non-stratified seeds.

To assess the effect of growth temperature, mechanically scarified EA seeds, which had been subjected to cold stratification, were sterilized and placed at RT or in a plant growth chamber maintained at 18 °C for 8 weeks. Germination rates of seeds grown at 18 °C were compared with those grown at RT.

### Experiment 2: GA_3_ treatments and water uptake

The effect of GA_3_ on seed germination was investigated using EA seeds. Mechanically scarified seeds, which had been subjected to cold stratification, were cultured on MS media supplemented with GA at concentrations of 0, 2, 5, or 10 mg/L. The petri-dishes were incubated in a plant growth chamber maintained at 18 °C for 8 weeks. Germination rates of seeds grown in the media containing GA were compared to those grown in the media without GA.

To determine how much water viola seeds absorb with time during germination, the total weight of the seeds was measured. Mechanically scarified EA seeds, which had been subjected to cold stratification, were sterilized and placed in a petri-dish lined with sterile Whatman filter paper. Seeds that did not undergo mechanical scarification were used as a control group, and water or 10 mg/L GA was added. The petri-dishes were placed in a growth chamber maintained at 18 °C for 4 weeks. After blotting the seeds with sterile Whatman filter papers to remove excess solution from the outer seed coats, the weight of the seeds was measured every 5 days. Change of seed weight was calculated based on the initial weight and compared to that of the control group.

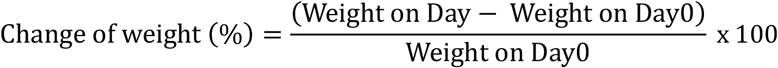

The seeds germinated at various stages were dissected and observed under a stereo microscope (MZ16F, Leica Microsystems, Wetzlar, Germany).

### Experiment 3: Agar concentration

As water uptake is considered a crucial factor for seed germination, the effect of agar concentration on the germination of viola seeds was investigated. Mechanically scarified EA seeds, which had been subjected to cold stratification, were sterilized and cultured on MS media containing agar (Sigma-Aldrich, St. Louis, MO, USA) at various concentrations of 0, 3, 5, or 7%. The seeds were placed in a plant growth chamber maintained at 18 °C. Germination rates were measured to determine the effect of different agar concentrations.

### Statistical analyses

The number of germinated seeds, defined as those that emerged either a root or a leaf, was counted weekly for 8 weeks. Each treatment included three replications. Data from each treatment were collected and compared to the control group using Tukey’s multiple range test (one-way ANOVA, P < 0.05) using JASP (Version 0.19.3; JASP Team, 2025).

## Results

A new method of mechanical scarification was developed to cut the tips of the seed coats, which facilitated the absorption of water and air without damaging the embryo (Figure 1). Soaking the seeds in water resulted in the soft seed coats, which enable to cut the seeds gently. Mechanically scarified EA seeds exhibited a germination rate of 10.67% on day 14, 26.67% on day 22, 36% on day 27, 61.33% on day 33, 63.33% on day 42, and 70% on day 54, whereas non-scarified seeds did not germinate (Figure 2). Initial germination, marked by the emergence of a root or a leaf, began within two weeks (Figure 3A and 3B). Following germination, leaf development was observed (Figure 3C), with roots emerging within five weeks (Figure 3D). While most seeds germinated within five weeks, some continued to germinate up to eight weeks.

**Figure 2.**
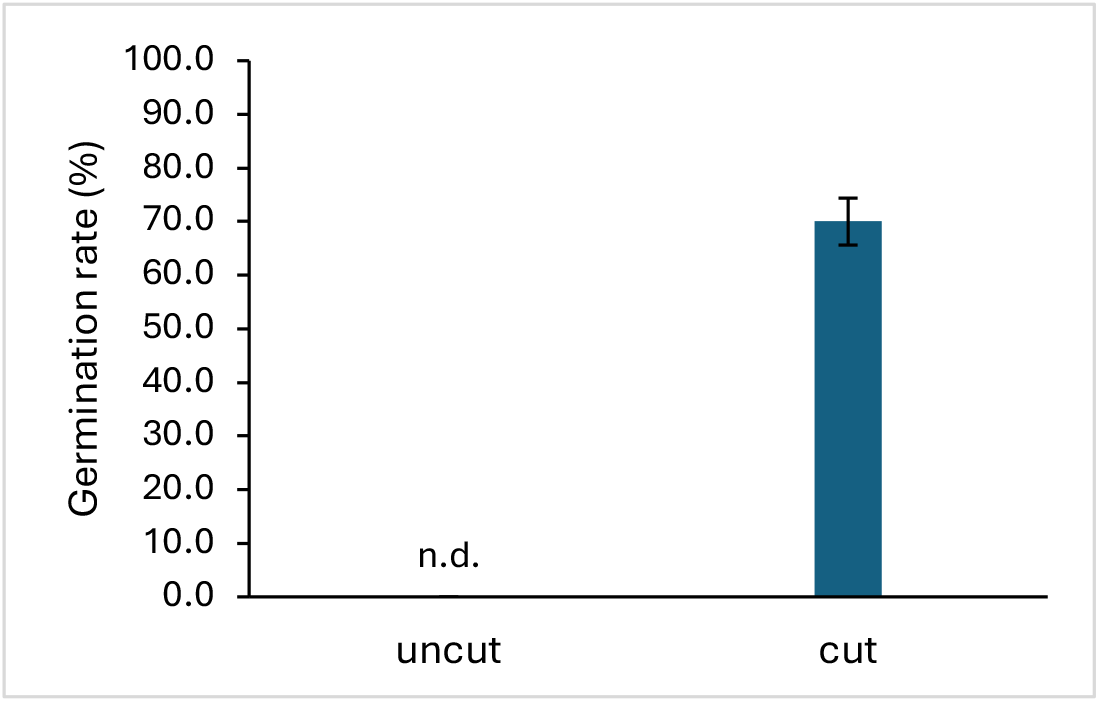
Germination rates of *Viola odorata* ‘Express Agusta’ seeds without and with mechanical scarification on Day 54. The seeds were treated cold stratification before the mechanical scarification. The bars indicate standard errors.

**Figure 3.**
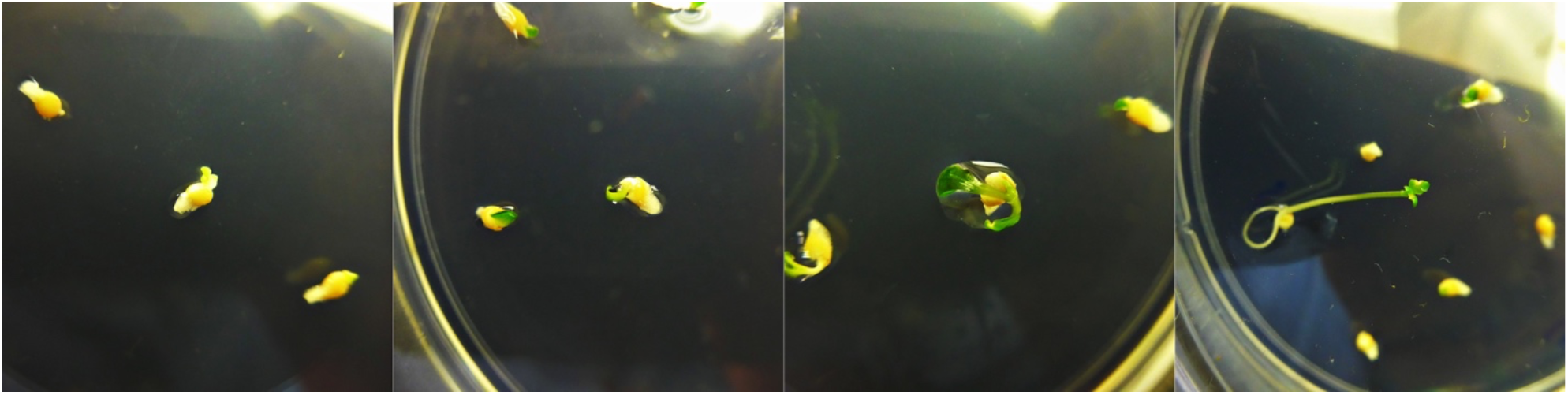
The developmental stages of *Viola odorata* ‘Express Augusta’ during seed germination. The seeds germinated (A) and emerged leaves (B) in two weeks. The leaves developed (C) and roots were shown (D) in five weeks.

Additional treatments were investigated to find optimal conditions for breaking seed dormancy. Cold stratification significantly enhanced the germination rate, approximately doubling the rate observed in EA seeds treated at RT (Figure 4A). The cold stratified seeds exhibited a germination rate of 70% on day 54, whereas non-stratified seeds showed a germination rate of 43.89% on day 54. A growth temperature of 18 °C was also proven effective, significantly increasing the viola germination rates (Figure 4B). The EA seeds cultivated at 18 °C demonstrated a germination rate of 70% on day 54, whereas the seeds grown at RT exhibited a 48.33% germination rate. The application of GA_3_ promoted both germination and subsequent plant growth. Germination rate in the media containing 2 and 5 mg/mL GA_3_ was not significantly different from the control group, but the rate increased significantly in media with 10 mg/mL GA_3_ compared to the media without GA_3_ (Figure 5).

**Figure 4.**
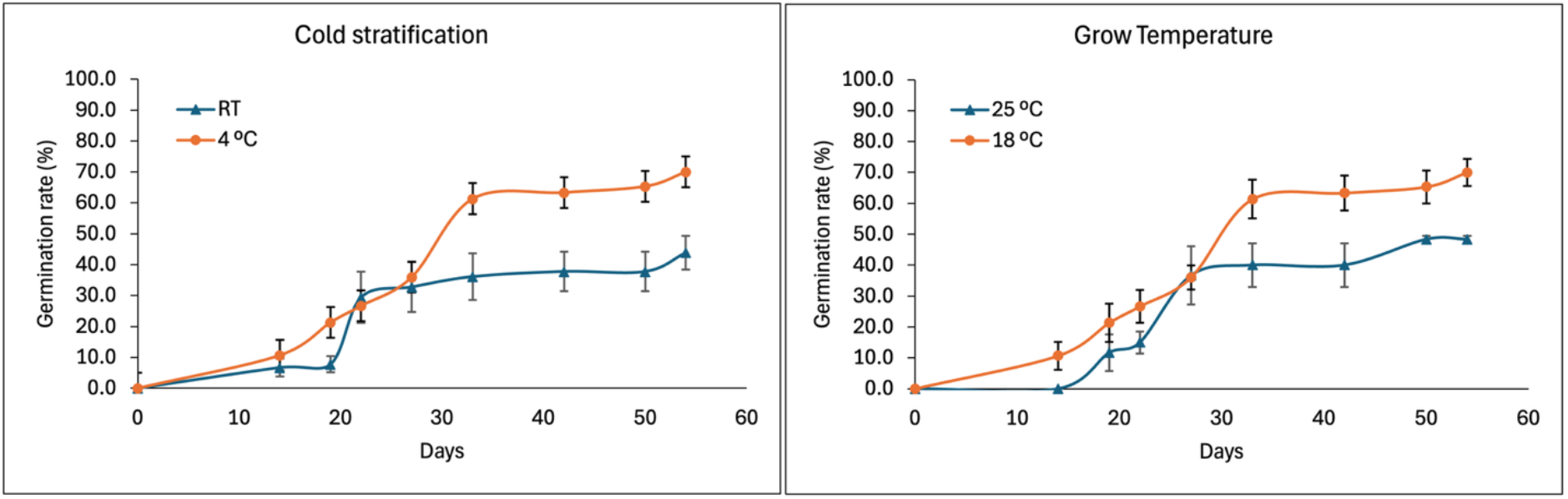
The ePect of cold stratification (Left) and growth temperature (Right) of *Viola odorata* ‘Express Agusta’ seed germination. The mechanically scarified seeds were stored at 4 ^°^C for cold stratification or room temperature (Left) and then placed at 18 ^°^C or room temperature (Right).

**Figure 5.**
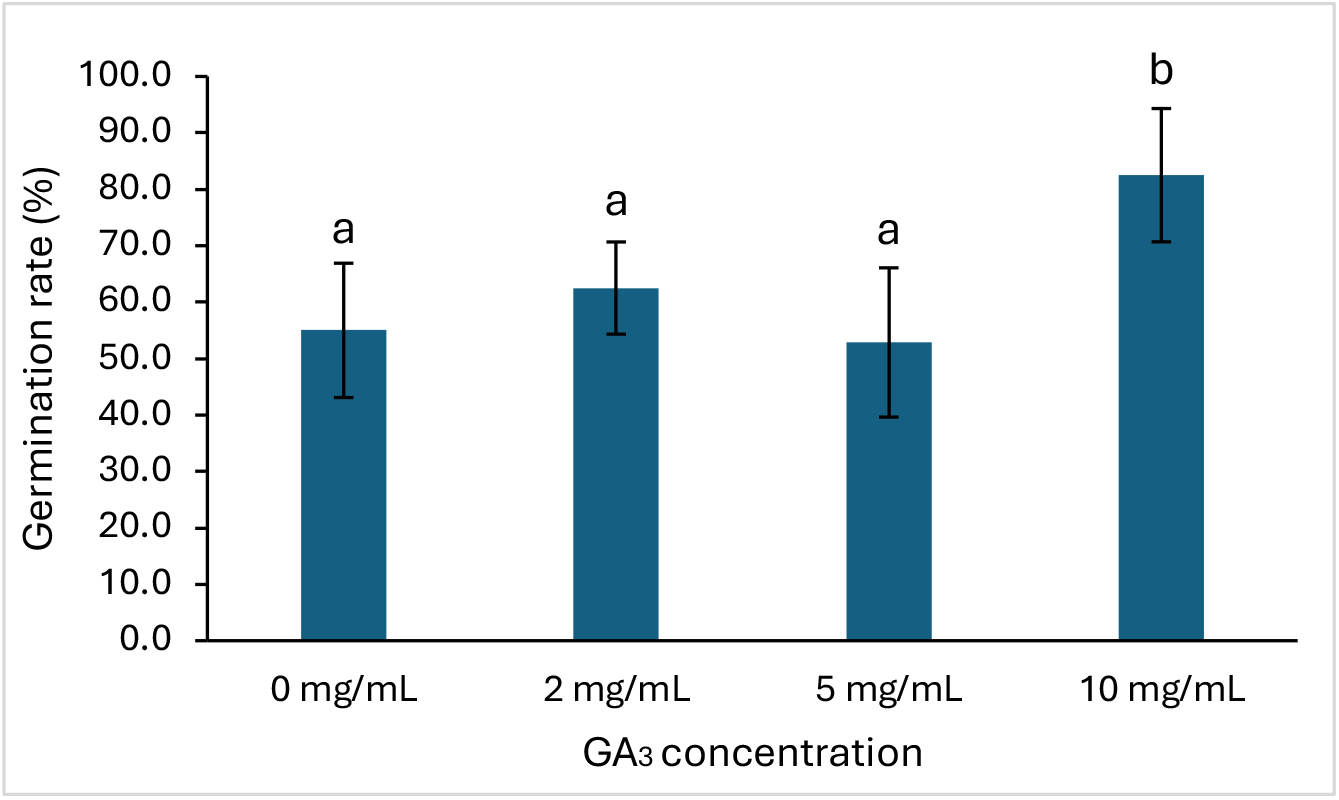
The ePect of GA on germination of *Viola odorata* ‘Express Augusta’ seeds on Day 54. After cold stratification, mechanically scarified seeds were grown in MS media containing various concentration of GA_3_ at 18 ^°^C. The bars indicate standard errors; the letters denote statistically significant differences between the treatments (p < 0.001, Tukey’s test).

The seeds dissected and observed under a microscope to investigate a change of embryo and endosperm during germination. The seeds not germinated didn’t exhibited any change inside (Figure 6A-1), and the embryo was bigger when started to germinate in two weeks (Figure 6A-2). The embryo was developed into roots or leave (Figure 6A-3 and 6A-4). To assess how much water viola seeds absorb during germination, seed weight was measured over time. Non-scarified seeds placed on a filter paper with water showed no significant change in weight. In contrast, mechanically scarified seed that was treated with both water and GA_3_ continuously increased in weight with time, and the seeds treated with GA_3_ exhibited a dramatic increase after day 15 (Figure 6B). The change of weight in the seeds treated with water was 1.5% on day 3, 3.1% on day 8, 3.9% on day 13, and 6.8% on day 20. The change of weight in the seeds treated with GA_3_ was 1.5% on day 3, 2.3% on day 11, 12.3% on day 18, and 15.9% on day 20. The application of GA_3_ resulted in a dramatic increase in weight. The effect of agar concentration on breaking seed dormancy was also investigated, but there was no significant difference in germination rates observed across concentrations of 0, 3, 5, and 7% (Table 1).

**Table 1.** The germination rates of *Viola odorata* ‘Express Augusta’ in MS media containing various concentration of agar.

|  | Day 0 | Day 14 | Day 22 | Day 27 | Day 33 | Day 42 | Day 50 | Day 54 |
| --- | --- | --- | --- | --- | --- | --- | --- | --- |
| 1% | 0.00 ± | 18.89 ± | 53.33 ± | 53.33 ± | 60.00 ± | 68.89 ± | 68.89 ± | 83.33 ± |
|  | 0.00* | 8.65 | 2.72 | 2.72 | 4.71 | 3.95 | 3.95 | 2.72 |
| 3% | 0.00 ± | 20.00 ± | 43.33 ± | 51.67 | 76.67 ± | 81.67 ± | 81.67 ± | 88.33 ± |
|  | 0.00 | 7.07 | 4.71 | ±1.18 | 16.50 | 12.96 | 12.96 | 8.25 |
| 5% | 0.00 ± | 2.22 ± | 26.67 ± | 33.33 ± | 56.67 ± | 68.89 ± | 68.89 ± | 84.44 ± |
|  | 0.00 | 1.72 | 13.66 | 13.66 | 9.31 | 5.24 | 5.24 | 8.48 |
| 7% | 0.00 ± | 11.11± | 46.67 ± | 51.11 ± | 71.11 ± | 74.44 ± | 74.44 ± | 77.78 ± |
|  | 0.00 | 3.75 | 6.83 | 3.75 | 10.99 | 8.48 | 8.48 | 9.47 |

\*The numbers indicate standard errors.

**Figure 6.**
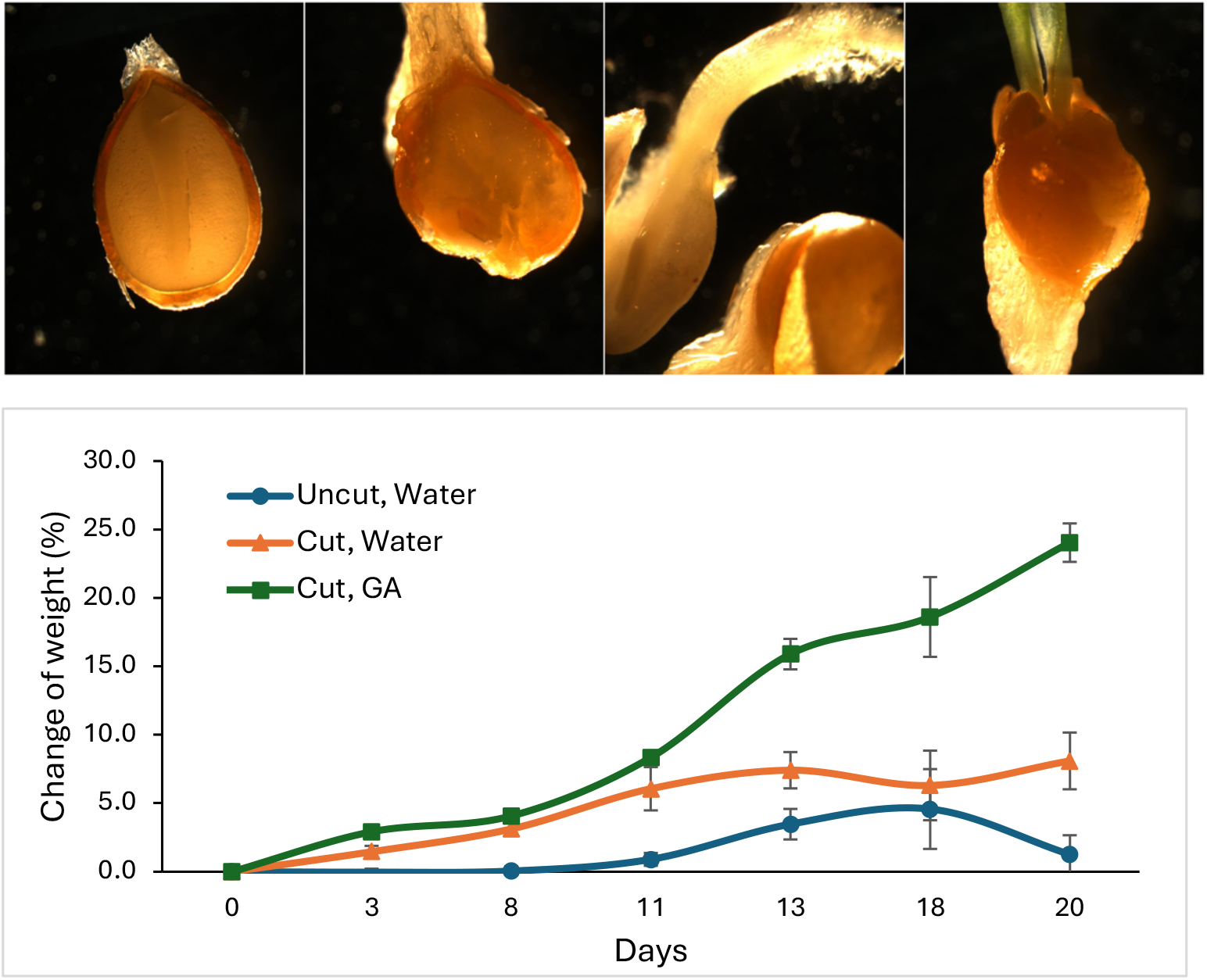
The change of seed weights with water or GA_3_ uptake in *Viola odorata* ‘Express Augusta’ seeds. After cold stratification, mechanically scarified seeds were grown at 18 C. The seed weights each day were compared to the weights on day 0. The bars indicate standard error.

## Discussions

Dormancy in *V. odorata* can be effectively broken through mechanical scarification. The hard seed coat of viola is a significant barrier to germination, and our experiments confirmed that non-scarified seeds of the EA variety did not germinate, which aligns with the finding of Barekat et al. (2013). A novel method of mechanical scarification was developed in this study, which involves creating a small, precise opening in the seed coats while minimizing harm to the embryo. This technique is an improvement over common methods, such as rubbing with sandpaper, which can often damage the embryo and lead to a lower germination rate. Mechanical scarifications enable seeds to absorb water. According to Finch-Savage and Leubner-Metzger (2006), germination rate is related to hydrotime and water potential. The weights of EA seeds increased over time by absorbing water in two weeks. The rapid increase in seed weight in EA seeds grown in GA_3_ solution observed after two weeks might be the effect of GA_3_ in accelerating plant growth, rather than just water uptake. GA_3_ is well known as an important hormone that plays a role in breaking seed dormancy and essential in plant growth by triggering cell division and the elongation process, transitions from meristem to shoot growth (Shah et al., 2023).

Nondeep physiological seed dormancy is regulated by the environmental cues activated through many different physiological mechanisms and temperature is a major factor (Finch-Savage and Leubner-Metzger, 2006). By combining with mechanical scarification, cold stratification increased EA germination. However, the effectiveness of stratification varies depending on the method and cultivars. For instance, cold stratification significantly increased the germination rate in this study, aligning with research by Shea and Colquhoun (2022) reported a germination rate of 68.8% for the ‘Rubra’ with an 8-week treatment. However, Barekat et al. (2013) reported a lower rate of 28.33% with 2-month cold treatment. In contrast, *V. pedata* exhibited improved seed germination in warm and dry conditions for 8 to 12 weeks, followed by stratification at 18 °C for 8 to 12 weeks after removing the elaiosomes (Gehring et al., 2013). In another case, *Rhus javanica* was successfully germinated through cold-moist stratification at 4 °C for 12 weeks after mechanical scarification (Cho et al., 2020). Optimal growth temperature and GA_3_ application are also beneficial in seed germination. EA seeds exhibited an enhanced germination rate when grown at 18 °C and when GA_3_ was provided.

The successful germination of other V. *odorata* varieties, including EA and RW, using the new mechanical scarification combined with cold stratification, suggests its potential for broad application to different varieties, provided optimal growth conditions of each variety are met. We suggest that mechanical scarification is the most effective method for breaking the dormancy of viola seeds.

## Data Availability Statement

The data presented in this study are available on request from the corresponding author.

## References

Barekat T, Otroshy M, Samsam-Zadeh B, Sadrarhami A, Mokhtari A. A novel approach for breaking seed dormancy and germination in Viola odorata (A medicinal plant). Journal of Novel Applied Sciences. 2013;2(10):513–6.

Baskin JM, Baskin CC. A classification system for seed dormancy. Seed science research. 2004 Mar;14(1):1–6.

Cho JS, Jang BK, Lee CH. Breaking combinational dormancy of Rhus javanica L. seeds in South Korea: Effect of mechanical scarification and cold-moist stratification. South African Journal of Botany. 2020 Sep 1;133:174–7.

Finch-Savage WE, Leubner-Metzger G. Seed dormancy and the control of germination. New phytologist. 2006 Aug;171(3):501–23.

Franklin S, Tran LB, Farzad D, Hill RI. Seed Germination In Viola pedunculata and Viola purpurea SUBSP. Quercetorum (Violaceae), Critical Food Plants For Two Rare Butterflies. Madroño. 2017 Jan 1:43–50.

Gehring JL, Cusac T, Shaw C, Timian A. Seed germination of Viola pedata, a key larval host of a rare butterfly. Native Plants Journal. 2013 Sep 21;14(3):205–12.

Górnik K, Sas-Paszt L, Seliga Ł, Pluta S, Derkowska E, Głuszek S, Sumorok B, Mosa WF. The effect of different stratification and scarification treatments on breaking the dormancy of Saskatoon berry seeds. Agronomy. 2023 Feb 11;13(2):520.

Gupta R, Chakrabarty SK. Gibberellic acid in plant: Still a mystery unresolved. Plant Signaling & Behavior. 2013 Jun 28;8(9):e25504. doi:10.4161/psb.25504

Keene SA. Laying the Groundwork: Developing Biochemical, Biotechnological, and Horticultural Tools for a Fragrance-Focused Breeding Program in Viola. University of Florida; 2022.

Keene, SA, Colquhoun, TA (2022). Germination of Viola odorata, a genetic resource for fragrance in Viola breeding. Combined Proceedings of the International Plant Propagators’ Society, 72, 127–132.

Keene SA, Sims M, Kim JY, Colquhoun TA. Temporal, Developmental, and Comparative Characterization of the Floral Volatile Emissions of the Famously Scented Violet Species, Viola odorata. HortScience. 2024 Jul 1;59(7):974–85.

Marcussen T. Allozymic variation in the widespread and cultivated Viola odorata (Violaceae) in western Eurasia. Botanical Journal of the Linnean Society. 2006 Aug 1;151(4):563–71.

Mittal, P., Gupta, V., Goswami, M., Thakur, N., & Bansal, P. (2015). Phytochemical and pharmacological potential of Viola odorata. International Journal of Pharmacognosy, 2(5), 215– 220.

Mousavi SR, Rezaei M, Mousavi A. A general overview on seed dormancy and methods of breaking it. Advances in environmental biology. 2011 Sep 1;5(10):3333–7.

Nawrot-Chorabik K, Osmenda M, Słowiński K, Latowski D, Tabor S, Woodward S. Stratification, scarification and application of phytohormones promote dormancy breaking and germination of pelleted scots pine (Pinus sylvestris L.) seeds. Forests. 2021 May 14;12(5):621.

Shah SH, Islam S, Mohammad F, Siddiqui MH. Gibberellic acid: a versatile regulator of plant growth, development and stress responses. Journal of Plant Growth Regulation. 2023 Dec;42(12):7352–73.

